# Interactive 3D visualization and post-processing analysis of vertex-based unstructured polyhedral meshes with ParaView

**DOI:** 10.1101/2021.10.15.464601

**Authors:** Paula C. Sanematsu

## Abstract

The development of physics-based 3D models that investigate the behavior of biological tissues requires effective and efficient visualization tools. The open-source software ParaView has such capabilities, but often impose a steep learning curve due to the use of the Visualization Toolkit (VTK) data structures. To overcome this, I show how to setup the components of 3D vertex-like models, i.e., vertices, faces, and polyhedra, into the VTK data format and then output as ParaView unstructured grid files. I present a few relevant tools to visualize and analyze the files in ParaView. All sample codes are available in the Github repository vis3Dvertex.

## 1. Introduction

The development of 2D continuum-[1], particle-[2], and vertex-based models [3-5] to understand the behavior of cellular tissues has revolutionized our understanding of how biological cells behave and interact with each other from a mechanistic point of view. One remarkable example is how vertex models allowed us to understand how epithelial tissue in the lungs behave differently for normal vs. asthmatic tissue [6]. In this work, the visualization of modeling and experimental results was crucial to understand the biological processes.

With the rapid advancement of biological imaging and computational power, it is reasonable to expect the further advancement of 3D vertex models as the 2D models rely on the assumption that a cross sectional plane of a 3D tissue is representative of the entire height of a monolayer tissue. Although this is a reasonable assumption in many instances, for various other cases, it is not [7, 8]. Beyond the monolayer configuration, researchers have developed 3D vertex models to understand how polyhedral-shaped cells behave in a three-dimensional tissue.

Studies as early as 2004 [9] developed 3D vertex models to understand cell deformation and rearrangement under external forces. Merkel and Manning [10] showed that a vertex-like 3D self-propelled Voronoi (SPV) model, governed by an energy functional depended on cell shapes exhibited a rigidity transition, similarly to the 2D vertex model. In general, vertex-like models in 2D and 3D include vertex [11] and Voronoi [10] models. The former has the cell vertices as the degrees of freedom whereas, in the latter, a Voronoi tessellation is created based on the cell centers which, in turn, are considered the degrees of freedom. Hereinafter, the term “3D vertex models” refers to the class of vertex-like models, including vertex and Voronoi models.

An essential component to the further advancement of 3D vertex models is the efficient visualization of simulation results. However, 3D visualization is not trivial because visualizing polyhedra requires rendering, that is, converting a 3D image into a 2D image in the computer. Rendering can be a computationally intensive task, which may limit the user’s possibilities while visualizing simulation results because every time the user changes the at angle, transparency, or coloring, a new rendering is performed. Thus, fast 3D rendering is indispensable for the visualization of 3D vertex models.

## 2. Problems and Background

In the published work of 3D vertex models, mostly two software have been used for visualization: POV-Ray [12] and MATLAB [13]. POV-Ray is a free and open-source ray tracing software that generates renderings based on a text-based scene description. It shows an intuitive representation of the data and has very high-resolution rendering, to the point that some renderings (not from vertex simulations) resemble real pictures. POV-Ray’s main disadvantage is the lack of user interaction. If the user wants to change the camera angle or the rendering color, those must be done in the text-based scene description file, and then re-render the visualization. MATLAB is a proprietary software with limited 3D rendering capabilities that includes camera angle changes, zooming in/out, but it lacks the ability to manipulate on the rendering.

Some scientific visualization software have been especially designed for fast 3D rendering of scientific data, such as VisIt [14], ParaView [15], and Avizo (Thermo Fisher Scientific). All have a GUI with a pipeline of input data and data manipulators rather than text-based interfaces like MATLAB or Matplotlib [16] that are commonly used for visualization of 2D simulations. Avizo is a commercial software widely used in the petroleum and geophysical communities. VisIt and ParaView are free and open-source and have extremely powerful parallelization capabilities. To put into perspective, the Department of Energy (DOE) Advanced Simulation and Computing Initiative (ASCI) developed VisIt for *terascale* simulations. ParaView was also designed to visualize and analyze extremely large datasets. It has successfully run on various platforms on 4000-32000 cores and it was able to visualize a billion-particle simulation [17]. Although parallelization of scientific visualization is not the focus of this work, ParaView allows this extension if parallelization of 3D vertex simulations becomes necessary.

In this work, I use ParaView to demonstrate how to visualize and analyze 3D vertex model simulations used in physics-based models. ParaView provides interactive visualization such that the user can view the 3D rendering from various angles, change color palettes, transparency, and rendering representation (e.g. wireframe, surface, volume) with a few mouse clicks. It contains filters that operate on the input data which can be manipulated, and then represented by plots, spreadsheets, or renderings. ParaView has an animation tool for time-lapse simulations to create movies or jump from a time step to another. Finally, it allows Python batch scripting without the need of using the pipeline.

ParaView handles its data structure using the Visualization Toolkit (VTK) [18], which may pose a steep learning curve for computational biologists, physicists, and engineers. To overcome such a hurdle, I briefly explain the VTK data structures necessary for a polyhedral mesh. I present a pseudocode to “convert” faces and vertices of polyhedral data into VTK data structures and output ParaView independent or a timeseries of files. I use the voro++ library [19] to create polyhedra by Voronoi tessellations. I modify voro++’s examples to create and output VTK data structures. All sample codes are available in vis3Dvertex along with a Singularity container image file (available on Github release page) that can be used to run the sample codes on a Linux machine with Singularity installed. In Section 4, I show how to visualize the output files in ParaView as well as how to manipulate the data using a few filters relevant for 3D vertex models. Although the focus of this work is on applications for biophysical models, this work is also relevant for any application that uses polyhedral unstructured meshes such as the materials science community who have used Voronoi-based models to understand material behavior under stress [20-23].

## 3. Paraview and VTK framework: Creating 3D polyhedral unstructured grids

### 3.1. VTK data structures and VTK polyhedral grids

I will briefly give some examples of VTK data structures and refer the reader to the free-to-download VTK user’s guide, textbook, and Doxygen manuals for more details: https://vtk.org/documentation/. The primary data structure in VTK is a data object. Data objects can be abstract such as graphs and trees or well-defined such as structured or unstructured grids – the latter being the focus of this work. In structured data, for example rectilinear grids, we know the connection between nodes (i.e. topology) and, therefore, we do not need to explicitly define the coordinates of each point. Unstructured data, on the other hand, require topology and point coordinates to be defined. Consequently, unstructured data demands considerably more memory, and one should only use it when structured grids are not possible.

A VTK structured or unstructured grid is composed of “cell types.” VTK supports various cell type dimensionalities such as vertex in 0D, line in 1D, triangle, quadrilateral, polygon in 2D, and tetrahedron, hexahedron, polyhedron in 3D (defined in the VTK source code `vtkCellType.h`). Cell types with a regular geometry, like tetrahedra (4 faces) and hexahedra (6 faces), use the vertices’ coordinates and a predefined ordering of the cell’s vertices to describe the cell topology. Thus, although we need to state the point coordinates, we do not need to explicitly define the topology of tetrahedral and hexahedral grids, saving some memory. In contrast, irregular polyhedral cells have a varying number of faces, and they need to have their topology explicitly defined along with their point coordinates. This work focuses on the polyhedral cells, represented by the `VTK_POLYHEDRON` cell type, to allow the visualization and analysis of the most general 3D unstructured grid that is used in physics-based 3D vertex models. Furthermore, the methodology presented here can be applied to experimental data whose vertex positions and topology are defined. Note that the `VTK_POLYHEDRON` only handles convex polyhedra; if concave polyhedra exist, then the `VTK_POLYGON` cell type can be used instead, such that a set of polygons would compose a polyhedron.

The topology or connectivity in polyhedron cells is stored as stream of ordered faces in the following format:

~~~
[numberOfCellFaces, (numberOfPointsOfFace0, pointId0, pointId1, …), (numberOfPointsOfFace1, pointId0, pointId1, …), …]
~~~

where `numberOfCellFaces` is the number of faces in the cell, `numberOfPointsOfFace0` is the number of points in the 0-th face, `pointId0` is the vertex index of point 0, `pointId1` is the vertex index of point 1 and so on.

Figure 1 shows one polyhedron and its face and vertex indexing lists from voro++’s modified example `cell_statistics_vtk.cc`.

**Figure 1:**
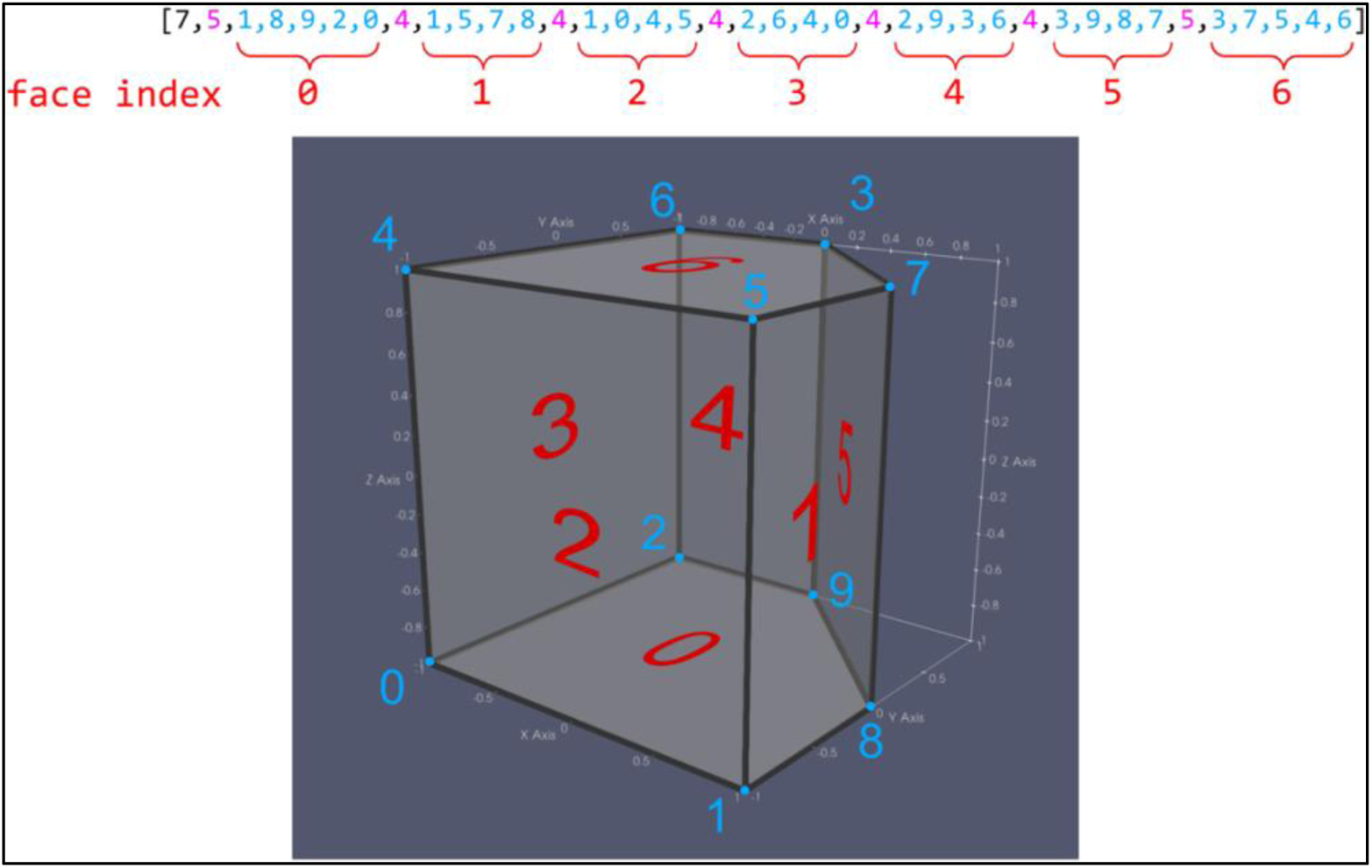
A polyhedron with labeled indices: vertex (blue) and face (red); and its `VTK_POLYHEDRON` face stream (top) created from voro++’s modified example `cell_statistics_vtk.cc`. The black number is the number of faces in the polyhedron, pink numbers are vertices per face, blue numbers are vertex indices, and red numbers are face indices.

To add a cell into the unstructured grid `vtkUnstructuredGrid`, I use the method `InsertNextCell`:

~~~
vtkIdType InsertNextCell(int cellType, vtkIdList *faceStream)
~~~

where `cellType` is `VTK_POLYHEDRON` and `faceStream` is shown in Figure 1.

The point coordinates are explicitly defined in the `vtkPoints` object and added to the `vtkUnstructuredGrid` with the method `InsertNextPoint:`

~~~
vtkIdType InsertNextPoint(double xCoordinate, double yCoordinate, double zCoordinate)
~~~

With cells and vertices defined, the basic components of an unstructured grid, I can now define attributes for the grid. Attributes can be variables used in the simulations such as time, pressure, velocity, force, surface area, volume, etc. These attributes are stored as data arrays whose number of components is defined by the user (see examples in Figure 2). Attributes can be point-, cell-, or field-based: `PointData` attributes are associated with the points whereas `CellData` attributes are associated with each polyhedron and assumed constant over the entire cell. `FieldData` gives a characteristic of the entire mesh – a common example is the time stamp.

**Figure 2:**
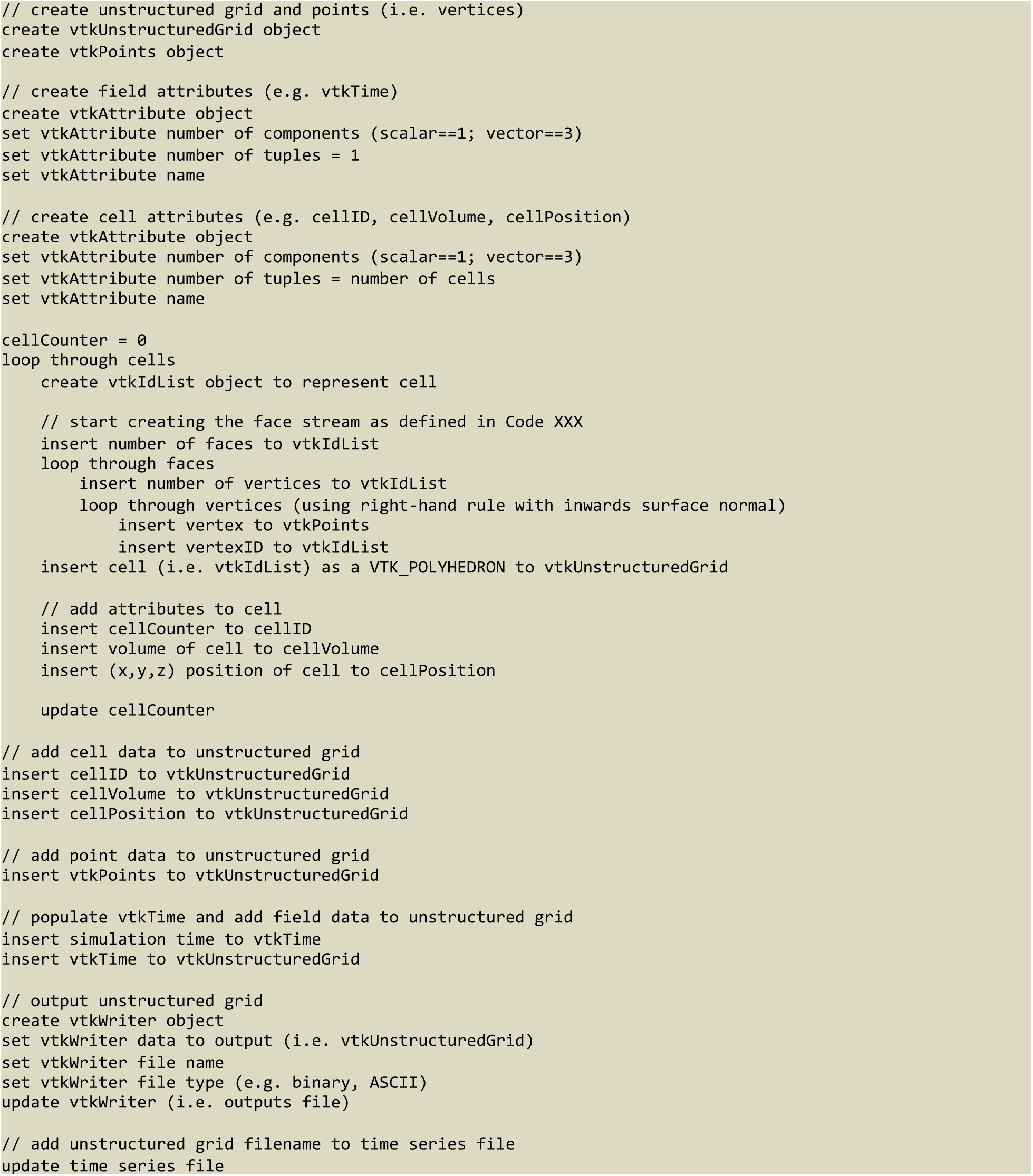
Pseudocode to create a polyhedral `vtkUnstructuredGrid` with `VTK_POLYHEDRON` cell type.

### 3.2. Pseudocode

Figure 2 provides a pseudo code of the concepts of Section 3.1. The first three blocks create the VTK objects for the unstructured grid and points objects. After these objects are created, three nested for-loops are necessary – cell, face, and vertex – to populate the `vtkPoints` object and to create the ID list of the `VTK_POLYHEDRON` cell type. In the vertex loop, I insert the points coordinate into `vtkPoints` and add the vertex index into the vtkIdList of `VTK_POLYHEDRON` (Figure 1, blue numbers). In the face loop, the number of vertices per face (Figure 1, pink numbers) are inserted into the `vtkIdList` of `VTK_POLYHEDRON`. After the face loop, I insert each cell attribute to its corresponding object.

After the nested for-loops, cell attributes objects (e.g. `cellID, cellVolume`) and are inserted into the unstructured grid as a `CellData` attribute. The points and their `PointData` attributes, if any, are also inserted into the unstructured grid. Finally, I output the unstructured grid using a `vtkWriter` object.

When the simulation iterates over time (or is minimized), one can write a ParaView timeseries file (.pvd) with the time stamp of each iteration and its corresponding “.vtu” unstructured grid file. For this iterative case, the pseudocode of Figure 2 would be contained within an iterative loop and each “.vtu” file needs the time stamp as a `FieldData` (second code block of Figure 2). Supplementary Figure S1 illustrates a timeseries for 5 iterations implemented in the voro++’s modified example `random_points_vtk.cc`.

### 3.3. Sample code using voro++

Figure 3 shows a snippet of `random_points_vtk.cc` with point coordinates insertion followed by the face loop where the `vtkFaces` object is populated for a single polyhedron. Note that in the voro++ library, the container that holds the Voronoi cells does not have a global list of vertices. The vertices are, instead, listed per cell. When two cells share a face with *N* vertices, these *N* vertices are listed twice in the global list. Thus, in the example `random_points_vtk.cc`, the global list of vertices, `points`, has repeated point coordinates. Other codes, however, may have a unique list of global vertices in which case the variable `containerVertexStartIndex` would not be necessary.

**Figure 3:**
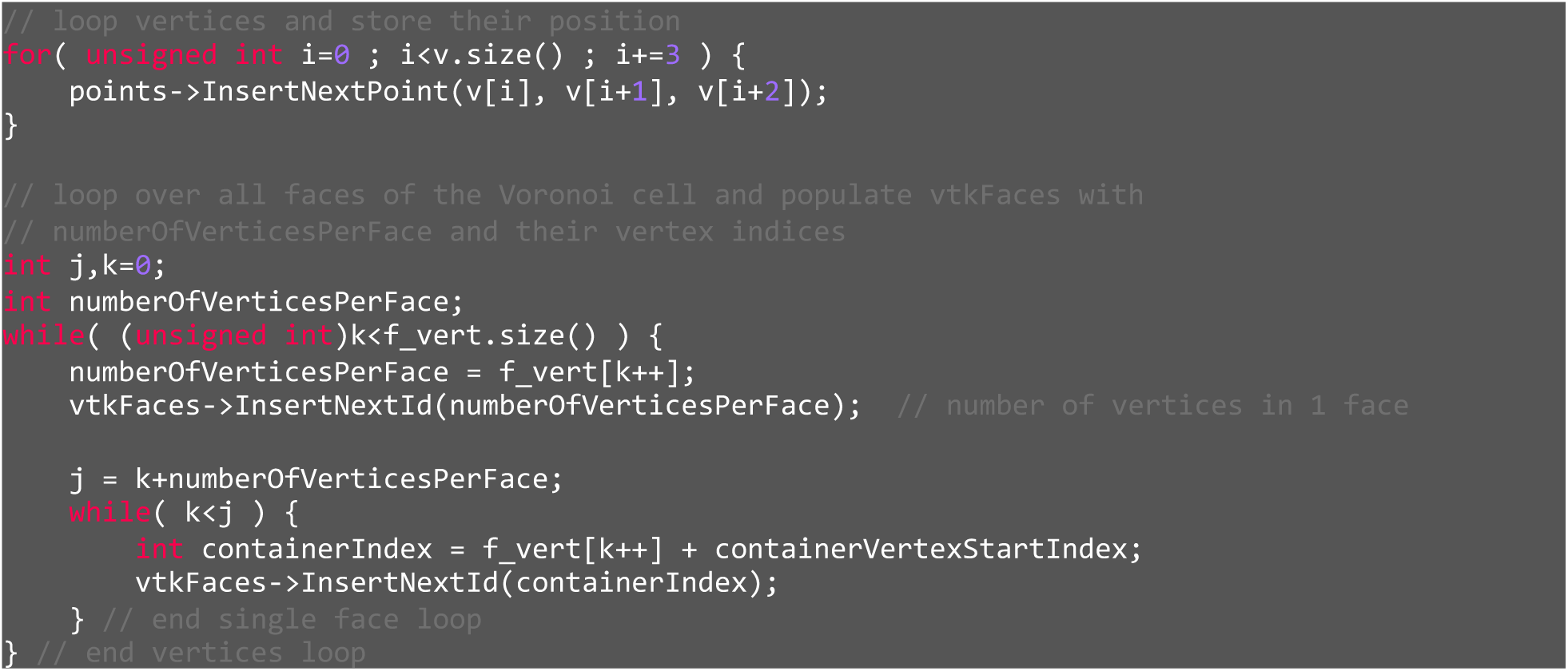
Snippet of point coordinate insertion and `VTK_POLYHEDRON` implemented in `random_points_vtk.cc`.

## 4. Implementation: ParaView basics

ParaView[15] works with visualization pipelines of sources, filters, and outputs. Figure 4 shows the main GUI components. In the “Pipeline Browser,” the user can view sources and filters along with their pipeline hierarchy indicated by the indentation. The user can select the “eye” on the left of the object to make it visible in the “3D

**Figure 4:**
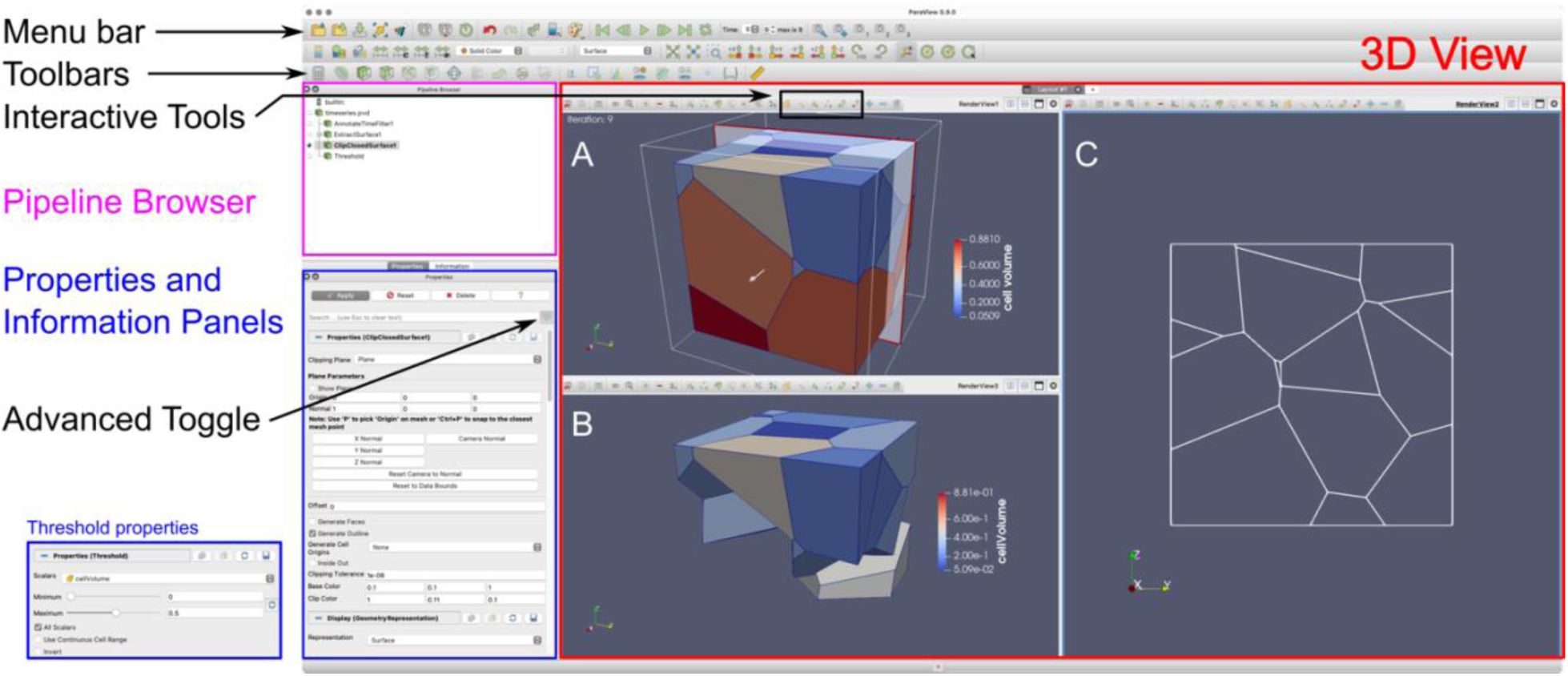
Paraview GUI. Figure adapted from Moreland [24] using voro++’s modified example `random_points_vtk.cc`. **(A)** The entire sample colored by cell volume. **(B)** After “Threshold” filter is applied with the criterion 0 ≤ cellVolume ≤ 0.5 (blue box). **(C)** A cross sectional plane at the plane indicated in panel A – for details, see SI Section 3.

View.” The “Properties” and “Information” panels are below the Pipeline Browser. These will display the properties and information of the pipeline selected object. The Properties panel also has the “Advanced Toggle” button which, if selected, displays additional properties about the object. Above the Pipeline Browser and 3D View, in the “Menu Bar,” the user can access most of ParaView’s features and “Toolbars,” which provides shortcuts to commonly used features. For an extended basic tutorial, refer to ParaView’s tutorial: The ParaView Tutorial version 5.4.1 [24], section – although an older version, the basics are mostly compatible with recent versions 5.9.X.

## 5. Illustrative examples: relevant filters and tools for 3D vertex models

All filters in ParaView are accessible through the Menu Bar (Filters -> Alphabetical) or through shortcuts in the Toolbar. The “Threshold” filter allows the user to define a scalar’s minimum and maximum threshold values. The cells within these limits will be displayed in the viewer. Figure 4 shows the entire Voronoi container before (panel A) and after the Threshold filter is applied (panel B and blue box).

The “Glyph” filter is useful to visualize vectorial data that can be displayed as a line to represent orientation or as an arrow that also includes the direction. In physics-based model, this representation is helpful to visualize velocity fields and cell orientation (polarity). Sahu, Schwarz [25] used the glyph filter to visualize cell stratification in the presence of heterotypic surface tension as shown in the blue-green-purple cells of Figure 5. The stratification becomes more evident with the cell orientation illustrated by the line glyphs positioned at the cell center. For more details on cell orientation, see Supplementary Information (SI) Section 2.

**Figure 5:**
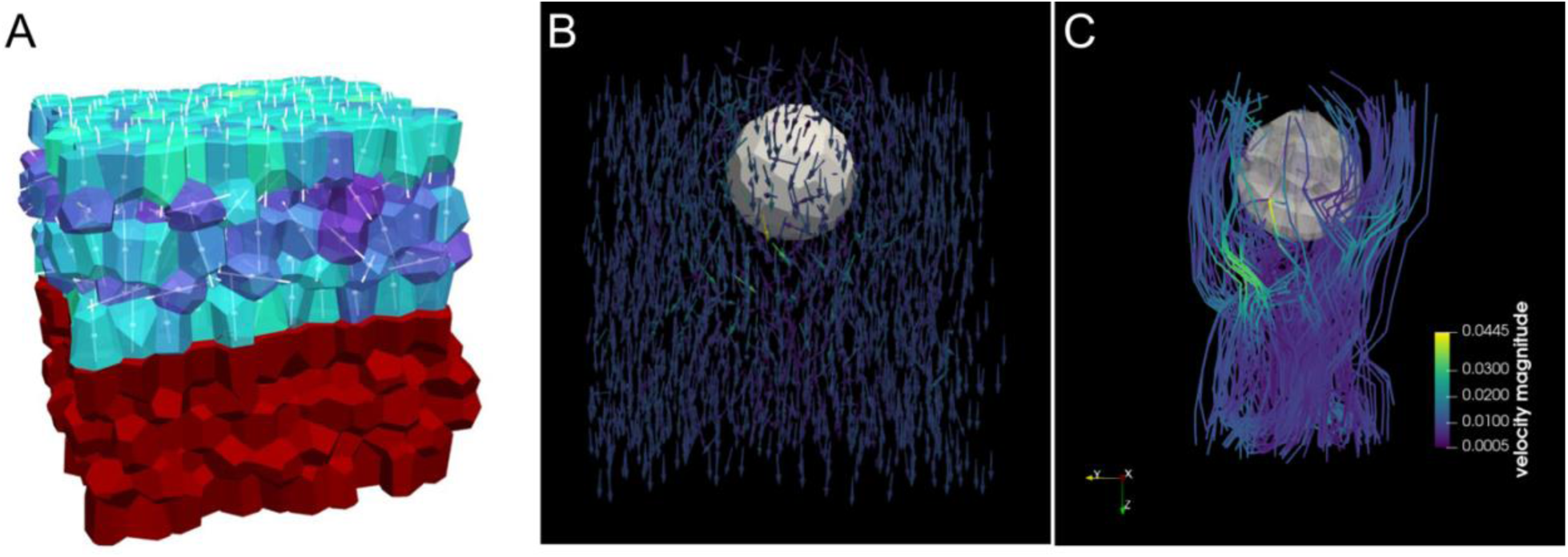
Glyph filter to show (A) cell orientation (reproduced from Sahu, Schwarz [25]; licensed under a Creative Commons Attribution (CC BY) license); (B) velocity field; and (C) Streamlines generated by Stream Tracer filter.

For simulations where cell velocity data is available, the filter “Stream Tracer” produces streamlines using a Runge-Kutta integrator on the velocity data. Here, to illustrate a meaningful example of 3D streamlines, I use a simulation from Sanematsu, Erdemci-Tandogan [26] to illustrate 3D streamlines around a spherical object as well the cells’ velocity field as arrow Glyphs.

The “Calculator” filter manipulates point or cell data by performing arithmetic operations. For cell-shape based models [10], Figure 6A shows how to calculate the cell shape parameter *s* = *S*/*V*^2/3^, where *S* is the observed cell surface area and *V* is the cell observed volume. This example shows how a filter can be used to derive data and reduce storage space.

**Figure 6:**
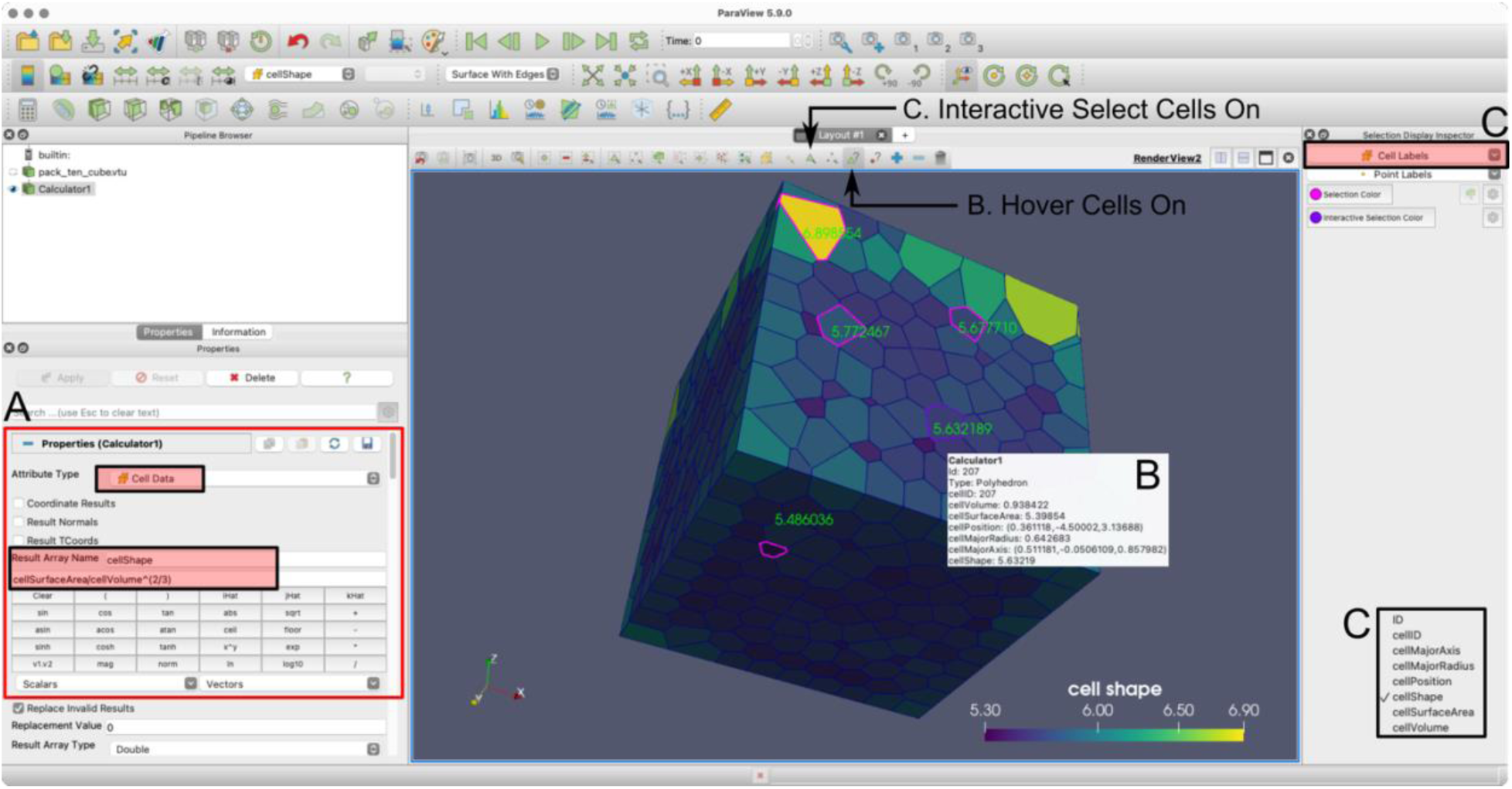
voro++’s modified example `import_vtk.cc.` (A) Calculation of cell shape parameter (*s* = *S*/*V*^2/3^) using the “Calculator” filter (red rectangle) to manipulate CellData. (B) Display of “Hover Cells On” of the purple outlined cell. (C) Pink outlined cells selected using “Interactive Select Cells On”: green numbers are the cellShape value that were selected by clicking on “Cell Labels” on the top right-hand corner.

In addition to filters, “Interactive tools” are very useful during development and development (Figure 6B, C). They display cell or point data as the user hover the mouse over cells. For implementation details refer to SI Section 4. Another practical feature is the “File -> Save State”, which saves the pipeline workflow in a “.pvsm” file. This state file can be later loaded (File -> Load State) and the pipeline workflow is applied to the original data or another dataset (see SI Section 5).

## 6. Conclusions

I present an efficient and powerful way to interactively visualize and analyze physics-based 3D vertex models using ParaView, an open-source software designed for scientific visualization of extremely large datasets. As ParaView uses the VTK library for its data structures, I first modify a very simple example from the voro++ library, `cell_statistics_vtk.cc,` to show how to “convert” a polyhedron’s vertices and faces into VTK data structures. I provide a general way to loop through a 3D-vertex model’s cells, faces, and points to create the VTK objects. I modify an example from the voro++ library, `random_points_vtk.cc`, to implement the pseudocode and create a timeseries file for time-evolving simulations. To visualize and analyze 3D vertex models, I present relevant ParaView filters for physics-based models by visualizing scalar and vectorial data. Other relevant tools that can be useful for debugging, such as the “Hovel Cells On,” are also presented. To generate such examples, codes are available in vis3Dvertex.

To start using ParaView can be a cumbersome task as the user has to become familiar with the pipeline workflow, VTK data structures, and polyhedral data structures. However, its existing capabilities of fast visualization, interactivity, and analysis are very useful to understand 3D vertex-models results in a timely manner. Here, I present examples to try to bridge the gap for biologists, biophysicists, engineers, and modelers so ParaView can be used to its potential. In addition, if it comes a day that 3D vertex models need CPU parallelization, ParaView is ready to be used.

## Acknowledgements

This work is supported by NIH grants R01GM117598 and R01HD099031. Simulations were performed on Syracuse University HTC Campus Grid, which is supported by NSF award ACI-1341006. The author thanks Dr. Lisa Manning’s mentorship and support for the realization of this manuscript, Dr. Larne Pekowski for patiently helping with Singularity and computing resources, and Dr. Preeti Sahu for providing valuable comments on this article.

## B-Required Metadata

**Table 1.** Code metadata

| Nr | Code metadata description |  |
| --- | --- | --- |
| C1 | Current Code version | V1.0.0 |
| C2 | Permanent link to code / repository used of this code version | https://github.com/pcsanematsu/vis3Dvertex |
| C3 | Legal Code License | GNU General Public License v2.0 |
| C4 | Code Versioning system used | git |
| C5 | Software Code Language used | c++ |
| C6 | Compilation requirements, Operating environments & dependencies | Required libraries: voro++, Eigen3, VTK; examples were tested on a Linux machine with the Singularity container image available on the release v1.0.0:<br>https://github.com/pcsanematsu/vis3Dvertex/releases/tag/v1.0.0. |
| C7 | If available Link to developer documentation / manual | https://github.com/pcsanematsu/vis3Dvertex#readme |
| C8 | Support email for questions |  |

