## Supplementary material for "Interactive 3D visualization and post-processing analysis of vertex-based unstructured polyhedral meshes with ParaView": Supplemetary Materials

Paula C. Sanematsu[1]
[1]: Department of Physics, Syracuse University, Syracuse, NY 13244, USA

### 1. *ParaView time series file*

```
<?xml version="1.0"?>
<VTKFile type="Collection" version="0.1"
         byte_order="LittleEndian"
         compressor="vtkZLibDataCompressor">
  <Collection>
    <DataSet timestep="0" group="" part="0" file="random_points_t_000.vtu"/>
    <DataSet timestep="1" group="" part="0" file="random_points_t_001.vtu"/>
    <DataSet timestep="2" group="" part="0" file="random_points_t_002.vtu"/>
    <DataSet timestep="3" group="" part="0" file="random_points_t_003.vtu"/>
    <DataSet timestep="4" group="" part="0" file="random_points_t_004.vtu"/>
  </Collection>
</VTKFile>
```

Figure S1: Example of a ParaView timeseries file (.pvd) that is implemented in the voro++'s modified example `random_points_vtk.cc`.

### 2. *Cell orientation – ellipsoid fit*

To obtain the cell orientation, I modify voro++'s example `voro++-0.4.6/examples/basic/import.cc` (`import_vtk.cc`) and include the code `fitEllipsoid.cc` in which an ellipsoid is fit in a polyhedron using the polyhedron's moment of inertia [1]. In `fitEllipsoid.cc`, the moment of inertia is calculated assuming each point has a unit mass. Note that there are other methods to calculate a polyhedron's orientation, such as considering it a uniform or heterogeneous solid. For simplicity and illustrative purposes, I use the point-mass moment of inertia which, as shown in [2], is suitable for some applications. For the full implementation of the point-mass moment of inertia, refer to the voro++'s modified example `import_vtk.cc` and its Glyph visualization is shown in Figure S2.

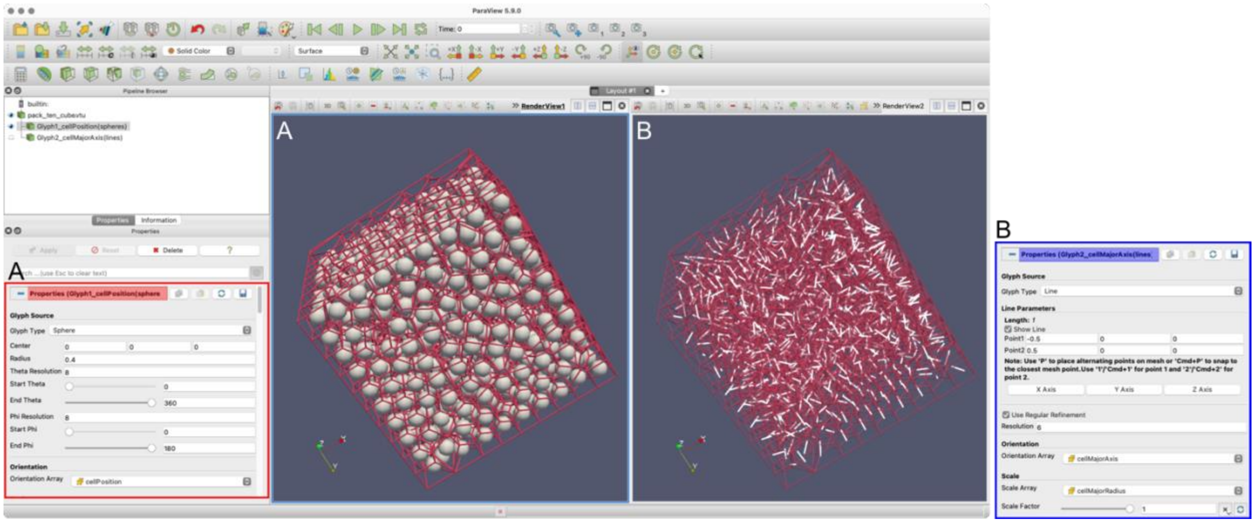


Figure S2: Glyph implementation to represent (A) cellPosition as spheres and (B) cellMajorAxis as lines. Red and blue panels show the properties of the glyph example.

#### 3. *2D cross sections*

Although imaging techniques for biological tissues have considerably improved in the past few years, the quantification of 3D cells shapes still poses difficulties. To overcome them, Sharp, Merkel [3] have demonstrated how to infer 3D cell shapes from 2D slices of an image stack. To reproduce such slices from 3D simulations, a sequence of filters in ParaView can be used: “ExtractSurface -> ClipClosedSurface” as displayed in Figure S3.

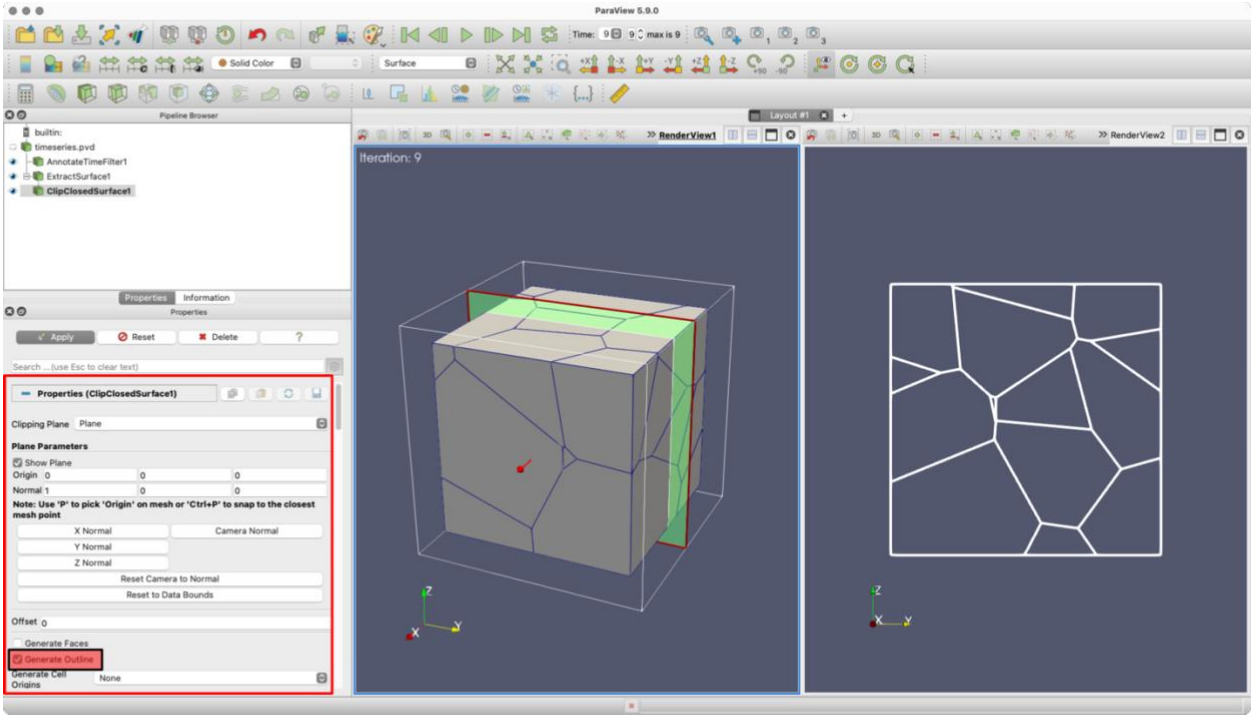


Figure S3: Extracting a 2D cross section of a polyhedral mesh using filters “Extract Surface -> ClipClosedSurface.” Check “Generate Outline” (small red box) in the “ClipClosedSurface” properties (big red box).

#### 4. *Ad-hoc cell and point information*

A few tools in ParaView are extremely useful during development and debugging stages. Here, I name them “Interactive tools” which are located right above the “3D View” as shown in **Error! Reference source not found.** (if not visible, go to “View -> Show Frame Decoration”). “Hover Cells On” displays the cell data as the user hover the mouse over cells (**Error! Reference source not found.**A) which can be done for point data with “Hover Points On.” **Error! Reference source not found.**B shows the “Interactive Select Cells On” tool, which selects specific cells (or points with the “Interactive Select Points On”) by mouse clicks and a cell data field is displayed next to the cell (to select which data field to be displayed, go to “View -> Selection Display Inspector”).

#### 5. *ParaView state files*

Two state files are included in the Github repository vis3Dvertex to reproduce Figures 4 and 6: “threshold_2Dcross_section_filters.pvsm” and “calculator_filter.pvsm”, respectively. To load “calculator_filter.pvsm”, go to “File -> Load State”, select “calculator_filter.pvsm” file, click “OK”. A “Load State Options” window will pop up (Figure S2). Ensure that “Choose File Names” is selected under “Load State Data File Options” and then click on the “…” to browse and find “pack_ten_cube.vtu” in your working machine (your path for the file name will look different than the Figure’s). For the state file “threshold_2Dcross_section_filters.pvsm”, choose the “timeseries.pvd” file.

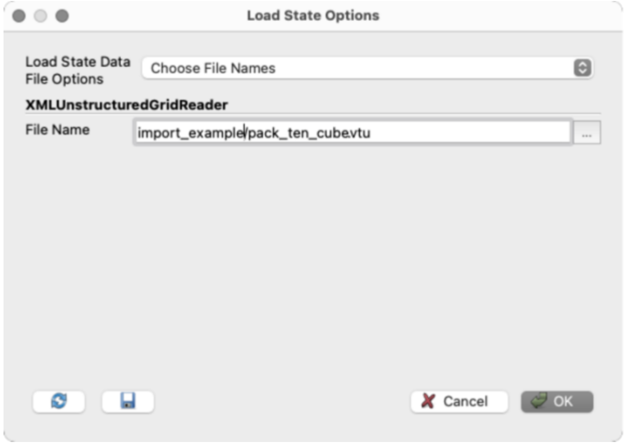


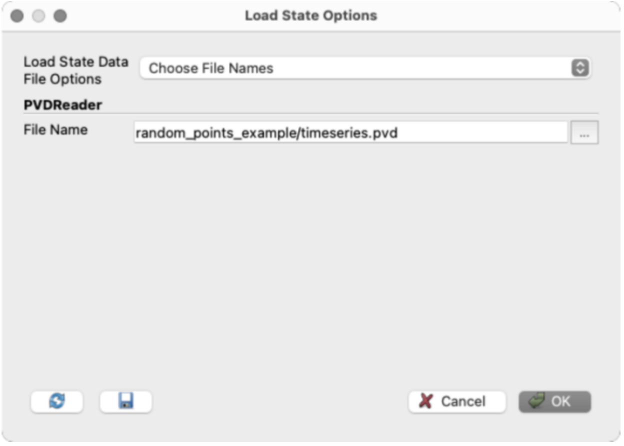


Figure S4: How to load a .pvsm file and select a specific .vtu file. Examples of how to select the appropriate files Figure 4 (left) and Figure 6 (right).

### *6. References*


1. Dobrovolskis, A.R., *Inertia of Any Polyhedron.* Icarus, 1996. **124**(2): p. 698-704.
2. Sahu, P., J. Schwartz, M., and M.L. Manning, *Geometric signatures of tissue surface tension in a three-dimensional model of confluent tissue.* arXiv preprint arXiv:2005.08429, 2021.
3. Sharp, T.A., et al., *Inferring statistical properties of 3D cell geometry from 2D slices.* PLoS One, 2019. **14**(2): p. e0209892.
